# Identification of a stress-sensitive endogenous opioid-containing neuronal population in the paranigral ventral tegmental area

**DOI:** 10.1101/2025.04.08.647881

**Authors:** Carrie Stine, David J. Marcus, Amanda L. Pasqualini, Ananya S. Achanta, Joseph C. Johnson, Sanjana Jadhav, Michael R. Bruchas

## Abstract

Nociceptin/orphanin FQ (N/OFQ), an endogenous opioid neuropeptide, and its G-protein coupled receptor NOPR have been implicated in motivation, feeding behaviors, and aversion. Stress-induced dysfunction in these states is central to the development of numerous psychiatric disorders, and the N/OFQ-NOPR system’s role in reward- and stress-related responses has driven broad interest in NOPR as a therapeutic target for anxiety and depression. However, the impact of stress on N/OFQ signaling in the context of its influence on discrete midbrain reward circuitry remains unknown. To this end, we focused on a possible candidate population of N/OFQ neurons in the paranigral ventral tegmental area (pnVTA^PNOC^) that have been shown to act locally on NOPR-containing VTA dopamine neurons to suppress motivation. Here we report and characterize pnVTA^PNOC^ sensitivity to stress exposure and identify a functional excitatory and inhibitory afferent input to this subpopulation from the lateral hypothalamus (LH). Our results indicate that pnVTA^PNOC^ neurons become recruited during exposure to a range of acute stressor types, whereas the GABAergic input from the LH to this population is suppressed by predator odor stress, providing a mechanism for disinhibition of these neurons. These findings suggest that this N/OFQ population in the pnVTA could act as a critical bridge between stress and motivation through interactions with upstream hypothalamic circuitry.

## INTRODUCTION

Stress exposure is a major risk factor in the development of addiction, relapse susceptibility, anxiety, and mood disorders, all of which collectively impose a staggering global health burden ^1,2,3,4^. While these disorders are vastly diverse, they all commonly involve the emergence of anhedonia and atypical motivation, indicating that the neurobiological mechanisms driving functional reward-related behaviors are highly susceptible to disruption in these disease states ^5,6,7,8,9^. Understanding the circuitry and neurobiological substrates central to reward processing that become altered by stress is a critical first step toward identifying viable therapeutic targets with improved function.

The mesolimbic pathway, comprised of dopaminergic projections from the ventral tegmental area (VTA) to the nucleus accumbens (NAc), plays a central role in processing and responding to reward ^10,11^. Converging animal ^12,13,14^ and human studies ^15,16,17,18^ have demonstrated that acute stress impacts neural activity within mesolimbic circuitry. In the VTA specifically, stress is generally found to have a net effect of suppressing VTA dopamine (DA) neuron activity ^19,20^. Recent studies have also expanded on the VTA’s molecular complexity by revealing diverse neuropeptide subpopulations, released both by the VTA itself and by upstream inputs such as the lateral hypothalamus (LH), that have significant influence over this critical reward circuitry ^21,22,23,24^.

Among these subpopulations, neurons in the paranigral nucleus of the VTA (pnVTA) enriched with the endogenous opioid peptide nociceptin/orphanin FQ (N/OFQ) have recently emerged as key regulators of motivated behavior ^25^. Our group previously reported that activation of these N/OFQ-expressing pnVTA neurons (pnVTA^PNOC^ neurons) suppresses motivated reward-seeking behavior and drives aversive responses. Notably, N/OFQ signaling through its cognate G-protein coupled receptor NOPR, which is largely expressed on VTA DA neurons ^26^, negatively regulates dopamine tone ^27^, paralleling the effects of stress. Despite widespread implications of N/OFQ in stress responses ^28^, whether stress impacts this particular pnVTA^PNOC^ population which is critically situated to regulate motivation and reward-related behaviors remains unexplored.

Here we employed *in vivo* calcium imaging with GCaMP to monitor pnVTA^PNOC^ neuronal dynamics during exposure to different modalities of acute stressors including foot shock, tail suspension, and predator-related stimuli. We also examined pnVTA^PNOC^ neuron engagement during exploratory behaviors with stressful components that promote risk avoidance or anxiety-like behavior. To further isolate the putative upstream circuitry underlying pnVTA^PNOC^ activation, we investigated synaptic inputs from the lateral hypothalamus (LH), a key region implicated in stress and motivation with known influence in the VTA on the expression of motivated behaviors ^29,30^. By combining optogenetics, electrophysiology, and behavioral assays, we aimed to uncover the connectivity and functional relevance of LH projections to pnVTA^PNOC^ neurons during stress exposure.

This study identifies the selective sensitivity of pnVTA^PNOC^ neurons to physical, environmental, and predatory forms of stress. Moreover, we identify the LH as a significant afferent input to pnVTA^PNOC^ neurons, offering a mechanistic basis for their engagement during stress. These findings contribute to a growing understanding of VTA circuitry in stress processing and identify a unique role of pnVTA N/OFQ neurons as a tenable bridge underlying stress regulation of motivated behavior.

## MATERIALS AND METHODS

### Animals

Adult (18–35 g) male and female *Pnoc*-IRES-Cre (PNOC-Cre), *Slc32a1*-IRES-Cre (VGAT-Cre), *Slc17a6*-IRES-Cre (VGLUT2-Cre), and VGLUT2-Flp x PNOC-Cre mice were group housed in the animal facility at 22–24 °C on a 12h/12h reverse light/dark cycle (9:00 AM lights off) in ventilated cages with ad libitum access to standard chow and water. All animals were monitored for health status daily and before experimentation for the entirety of the study. Animal procedures were approved by the Animal Care and Use Committee of the University of Washington and conformed to US National Institutes of Health guidelines. All resources are listed in **Table S1**.

### Stereotaxic surgery

All coordinates, viruses, and volumes for experiments are listed in **Table S2**. After acclimating to the holding facility for at least seven days, mice were anesthetized in an induction chamber (1%-4% isoflurane) and placed into a stereotaxic frame (Kopf Instruments, model 1900) where they were maintained at 1%-2% isoflurane. A blunt needle syringe (86200, Hamilton Company) was used to deliver virus at a rate of 100nL/min either in the LH or pnVTA. For mice receiving intracranial fiber photometry implants, an optic fiber (400µm core, 2.5mm ferrule, Doric) was slowly lowered to 0.05mm above the injection site and secured using MetaBond (C & B Metabond). A stainless-steel headring was also secured on animals undergoing air puff to allow for head-fixation. Animals were allowed to recover from surgery for a minimum of 3 weeks before any behavioral testing, permitting optimal viral expression.

### Fiber photometry recordings

Fiber photometry studies were completed as described previously ^31^ (see **Supplementary Methods**). In brief, GCaMP6s fluorescence was excited using a 470 nm LED (Ca^2+^-dependent signal) and a 405 nm LED (isosbestic control, Ca^2+^-independent signal). LED intensities were set to 30µW at the optic fiber tip. GCaMP6s emissions were filtered (525 ± 25 nm), detected with a photoreceiver, and recorded by a real-time processor. For ChrimsonR stimulation, a 635 nm laser delivered 1mW red light at the fiber tip ^32^.

### In vivo animal experiments

All animal behaviors were performed within a sound-attenuated room maintained at 23°C at least one week after habituation to the holding room. Animals were handled for a minimum of three days prior to experimentation and were habituated to fiber photometry patch cord attachment to their fiber implants. For all experiments, mice were brought into the experimental room and allowed to acclimate to the space for at least 30 minutes prior to any testing. All experiments were conducted in red light to accommodate the reverse light cycle schedule, unless otherwise stated. All sessions were video recorded.

### Behaviors

#### Cued foot shock

Mice were placed in Med Associates Fear Conditioning Chambers (NIR-022MD) which consisted of a 29.5 x 23.5 x 21 cm chamber with a conductive grid floor lit by infrared light and contained within a soundproof box. Mice were exposed to ten 10s tones co-terminating with a 2s 0.5 mA shock with a variable inter-trial interval (VITI) of 45s-90s.

#### Tail lift

Mice were placed in a 10” x 10” clear acrylic box illuminated by a dim, diffuse white light 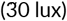. Mice were suspended by the tail four times for 20s with a VITI of 120s-180s. All suspensions were made to the same height.

#### Air puff

Four days prior to testing, mice were habituated to head-fixation on the OHRBETS platform as described previously ^31,33^. A fixed, solenoid-controlled O_2_ valve was positioned above the animal’s left whiskers. Mice were exposed to fifteen 0.1s 20 PSI air puffs with a VITI of 45s-75s. Solenoid opening (Parker, 003-0257-900) was controlled using an Arduino Mega 2560 REV3 (Arduino) and custom Arduino programs.

#### Looming

Mice were placed in a white-walled plexiglass arena (50 x 50 cm) illuminated by a diffuse white light 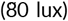 and allowed to roam freely. Looming was simulated four times via a posterboard blocking the arena’s overhead lighting (arena illumination reduced to 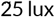) for 1s-2s, with a VITI of 120s.

#### Odor delivery

Mice were placed in a polyethylene chamber approximately 12 x 12 x 24 cm where air was continuously vacuumed out at a rate of 2L/min ^34^. To minimize odor release into the room, the chamber was placed in a fume hood and vacuumed air was passed through a carbon filter. Odors (2% 2MT or 2% peppermint oil, in separate sessions) were delivered four times per session for 30s periods with a VITI of 120s-180s.

#### Tail suspension test (TST)

TST was completed as described previously ^35^. Following a 5-minute baseline in a 10” x 10” clear acrylic box, mice were suspended by the tail continuously for 6 minutes. On counterbalanced test days, mice received either 0 or 20 Hz (5ms pulse width, 1mW laser power) cycling ON for 5s and OFF for 15s throughout the suspension (18 total cycles).

#### Open field test (OFT)

OFT was completed as described previously ^34,36^ in a white-walled plexiglass arena (50 x 50 cm) illuminated by a white light 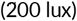. Center zone was defined as the middle 50% of the arena size. Mice were allowed to roam the arena freely for 30 minutes.

#### Elevated zero maze (EZM)

EZM was completed as described previously ^34,36^ in a circular maze (Harvard Apparatus) with a 200 cm circumference comprise of four 50 cm sections (two open and two closed ‘arms’), elevated 50 cm above the floor illuminated by a white light 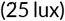 The maze path was 4 cm wide with a 0.5 cm lip on each open arm and 17 cm walls on each closed arm. Mice were positioned head-first into a closed arm and allowed to roam the maze freely for 7 minutes.

### Electrophysiology and ex vivo optogenetics

Coronal brain slices for electrophysiology recordings were prepared as previously described ^37^ (see **Supplementary Methods**). LH GABAergic and glutamatergic input onto pnVTA^PNOC^ neurons was evaluated using voltage clamp recordings of current responses in mCherry-labeled neurons following ChR2 excitation across a range of intensities.

### Tissue preparation and immunohistochemistry (IHC)

Unless otherwise stated, animals were transcardially perfused with 0.1M phosphate-buffered saline (PBS) followed by 40mL 4% paraformaldehyde. Brains were dissected and post-fixed in 4% paraformaldehyde overnight and then transferred to 30% sucrose solution for cryoprotection. Brains were sectioned at 30µm on a microtome and stored in 0.1M phosphate buffer at 4°C prior to immunohistochemistry and tracing experiments. Immunohistochemistry was completed as described previously ^31,38,39^ (see **Supplementary Methods**). For behavioral cohorts, viral expression and optic fiber placements were evaluated before inclusion in the presented datasets.

### Fiber photometry analysis

Fiber photometry data were analyzed as described previously ^31^. In brief, custom MATLAB scripts were used to normalize signal by detrending decay from bleaching then dividing by an LLS fit of the isosbestic trace scaled to the signal. The processed traces were extracted in windows surrounding the onset of relevant behavioral events (tail lift, odor, shock, air puff, looming, open arm entry, center entry), z-scored relative to the mean and standard deviation of each event window, and then averaged.

### Statistical analyses

Statistical analyses were performed as indicated in GraphPad Prism 9 and MATLAB 9.9 (MathWorks). All data are expressed as mean ± SEM unless specified otherwise.

## RESULTS

### pnVTA^PNOC^ neurons exhibit sustained activity throughout acute stress exposure across multiple stressors

N/OFQ-containing neurons in the pnVTA (pnVTA^PNOC^ neurons) act to suppress motivated behaviors. Stress is also known to disrupt motivation, but despite evidence linking N/OFQ with stress, the impact of stress on the activity of this population remains unknown. Importantly, the effects of stress exposure on motivation can vary depending on the form and duration of the stressor. To evaluate the effects of diverse stress conditions on pnVTA^PNOC^ activity, we injected PNOC-Cre mice with a Cre-dependent GCaMP6s (AAV-DJ-Ef1a-DIO-GCaMP6s) and implanted optic fibers in the paranigral ventral tegmental area (pnVTA) to record the calcium activity of pnVTA N/OFQ neurons (pnVTA^PNOC^) during exposure to a variety of stressful stimuli (**Figure 1A, B**). At 3-4 weeks post injection we detected robust, transient activation of pnVTA^PNOC^ neurons in response to a mild foot shock (**Figure 1C–F**, 1-way ANOVA F_3,24_=14.26, p<0.0001), the onset and offset of a 20-second tail lift (**Figure 1G–I**, 1-way ANOVA F_4,44_=10.94, p<0.0001), and a brief 0.1s air puff delivered to the whiskers (**Figure 1J–L**, 1-way ANOVA F_2,4_=5.417, p=0.0727). Interestingly, pnVTA^PNOC^ activity was time-locked with the duration of the stressor, returning quickly to baseline levels following the 0.1s air puff or 2s foot shock, but remaining elevated throughout the 20s tail suspension (**Figure 1E, H, K**). Across these stressors we observed similar activity in both male and female mice, although foot shock elicited a larger response in females (**Supplementary Figure 1A,B**). Notably, pnVTA^PNOC^ neurons were not activated in response to the 10-second tone that preceded each foot shock (**Figure 1E,F**, Tukey’s test baseline vs. tone p=0.9993), suggesting that pnVTA^PNOC^ neurons have selective sensitivity to stress rather than simply responding indiscriminately to any salient stimuli.

**FIGURE 1.**
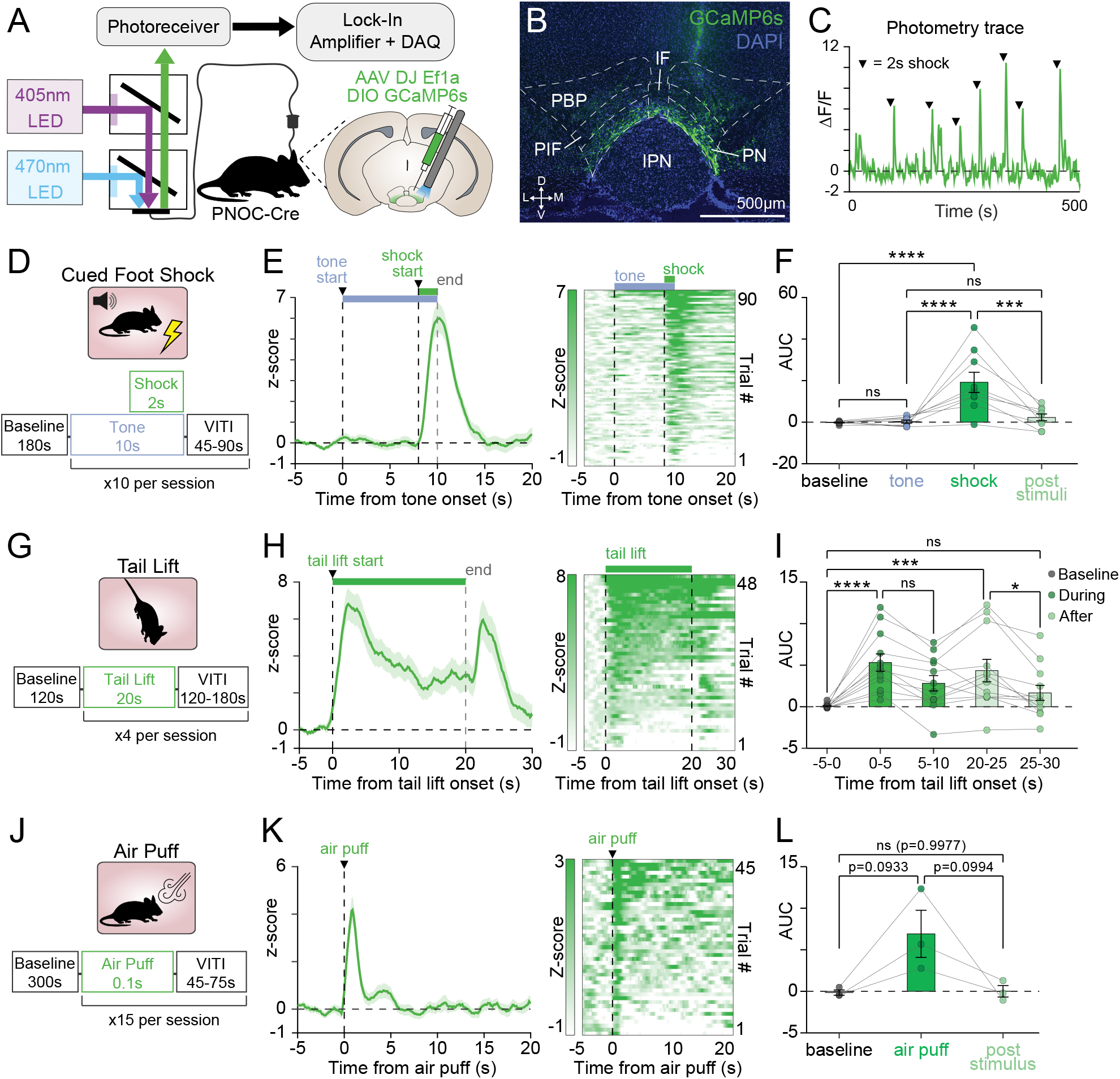
pnVTA^PNOC^ neurons are activated during exposure to acute stressors. **A** Fiber photometry schematic and cartoon of DIO-GCaMP6s (GCaMP6s) viral injection and fiber implant in the pnVTA of PNOC-Cre mice. **B** Representative coronal image showing DAPI (blue) and GCaMP6s (green) expression in pnVTA. **C** Representative trace of GCaMP6s F/F fluorescence throughout a cued foot shock session. Black arrows are aligned with foot shock onset. **D** Trial structure for a cued foot shock session (10s tone co-terminating with 2s 0.5 mA shock). **E** Left: Averaged trace of pnVTA^PNOC^ GCaMP6s activity during epoch surrounding tone-cued foot shock, aligned to tone onset. Right: Heat map of GCaMP6s fluorescence during same epoch, each row correspond to a trial in the averaged trace (left). (N = 9 mice). **F** Area under the curve (AUC) for averaged traces from E, calculated over 8-second intervals surrounding cued-foot shock events. GCaMP6s signal increases in response to shock but not tone (1-way ANOVA with Tukey’s multiple comparisons test, ****p < 0.0001, ***p < 0.001, N = 9 mice). **G–I** Same as D–F but for pnVTA^PNOC^ GCaMP6s activity during 20s tail lift. Activity averaged in 5-second intervals surrounding each tail lift shows increases first during tail lift and again when animal is lowered to the ground (1-way ANOVA with Tukey’s multiple comparisons test, ****p < 0.0001, ***p < 0.001, *p < 0.05, N = 12 mice). **J–L** Same as D–F but for pnVTA^PNOC^ GCaMP6s activity during acute air puff (0.1s, 20 PSI). Activity averaged in 5-second intervals surrounding each air puff (1-way ANOVA with Tukey’s multiple comparisons test, N = 3 mice). All data represented as mean ± SEM.

We next evaluated pnVTA^PNOC^ dynamics during innately anxiogenic exploratory behaviors (**Figure 2A**). In the open field test (OFT), we detected a significant increase in calcium activity as animals transitioned from the ‘safe’ edge of the arena to the open, ‘risky’ center (**Figure 2B, D**, 1-way ANOVA F_2,30_=16.18, p<0.0001). We observed a similar increase in the elevated zero maze (EZM) as animals entered the unprotected open arms of the maze (**Figure 2C, E**, 1-way ANOVA F_2,30_=6.305, p=0.0052). No sex-dependent effects were identified in either OFT or EZM (**Supplementary Figure 1C,D**). These findings demonstrate that pnVTA^PNOC^ neurons are also engaged by innately stressful environmental stimuli. Taken together, our results indicate that multiple forms of acute stress can elicit robust activation of N/OFQ-containing pnVTA neurons.

**FIGURE 2.**
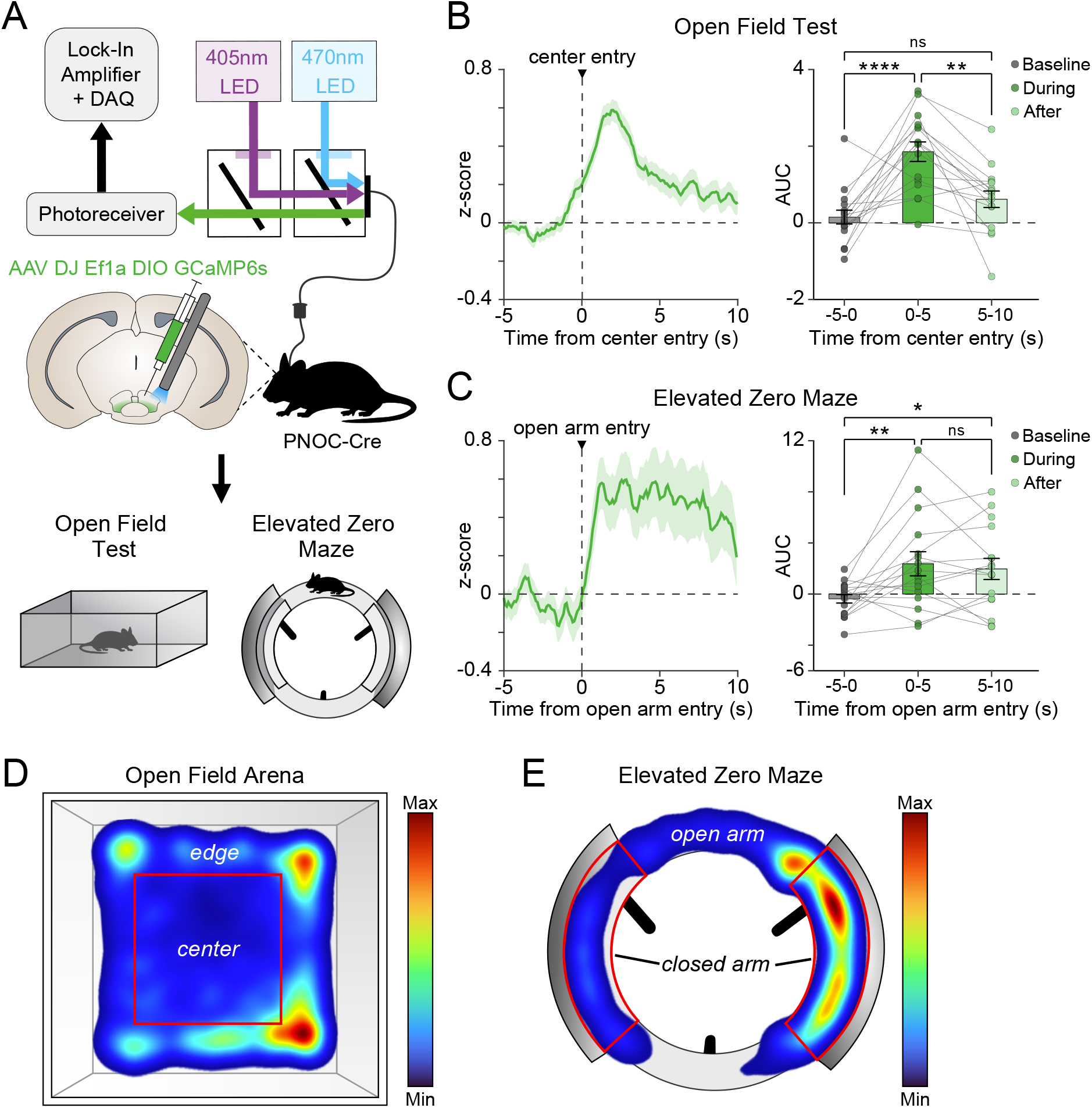
Anxiogenic exploratory behaviors drive pnVTA^PNOC^ activity. **A** Cartoon of DIO-GCaMP6s (GCaMP6s) injection and fiber implant into pnVTA of PNOC-Cre mice. GCaMP6s activity was recorded during open field test (OFT) and elevated zero maze (EZM). **B** Left: Averaged traces of pnVTA^PNOC^ GCaMP6s activity during high-anxiety epochs of the OFT, aligned to entries into the center zone of the open field arena. Right: Area under the curve (AUC) for averaged traces (left) calculated over 5-second intervals surrounding center zone entry. GCaMP6s activity increases during and immediately after center entry (1-way ANOVA with Tukey’s multiple comparisons test, ****p < 0.0001, **p < 0.01, N = 16 mice). **C** Same as B but for pnVTA^PNOC^ GCaMP6s activity during high-anxiety epochs of the EZM, aligned to entries into either open arm of the maze (1-way ANOVA with Tukey’s multiple comparisons test, **p < 0.01, *p < 0.05, N = 16 mice). **D** Heat map from a representative animal showing proportion of time spent in each area of the open field arena. Heat map shows more time spent around the edge than in the center. **E** Same as D but showing proportion of time spent in each area of the elevated zero maze. Heat map shows more time spent in the closed arms than in the open arms. All data represented as mean ± SEM.

### Predator odor stress engages pnVTA^PNOC^ neurons in both sexes, while predator looming stress elicits responses only in males

We further characterized the stress sensitivity of pnVTA^PNOC^ neurons by recording calcium activity following different predatory stressors (**Figure 3A**). When exposed to a looming stimulus that mimics the threat of an overhead predator, male mice displayed a significant increase in pnVTA^PNOC^ calcium activity while females did not (**Figure 3B,C**, 2-way ANOVA males vs. females F_1,10_=11.14, p=0.0075). We also assessed pnVTA^PNOC^ calcium activity dynamics in response to an aversive predator odor (2% 2MT, a predator urine derivative ^40^) and a non-aversive novel odor (2% peppermint oil), which served as a control for salience (**Figure 3D**). The aversive 2MT predator odor evoked a robust, sustained increase in calcium activity, whereas the non-aversive peppermint odor elicited no response (**Figure 3E**, 2MT: 2-way ANOVA F_4,32_=5.713, p=0.0014; peppermint oil: 2-way ANOVA F_4,32_=0.3368, p=0.8511). These data are consistent with our initial findings that pnVTA^PNOC^ neurons are selectively activated by stressful stimuli, and reveal a potential sex-dependent circuit level effect in response to certain predatory stressors, in male mice.

**FIGURE 3.**
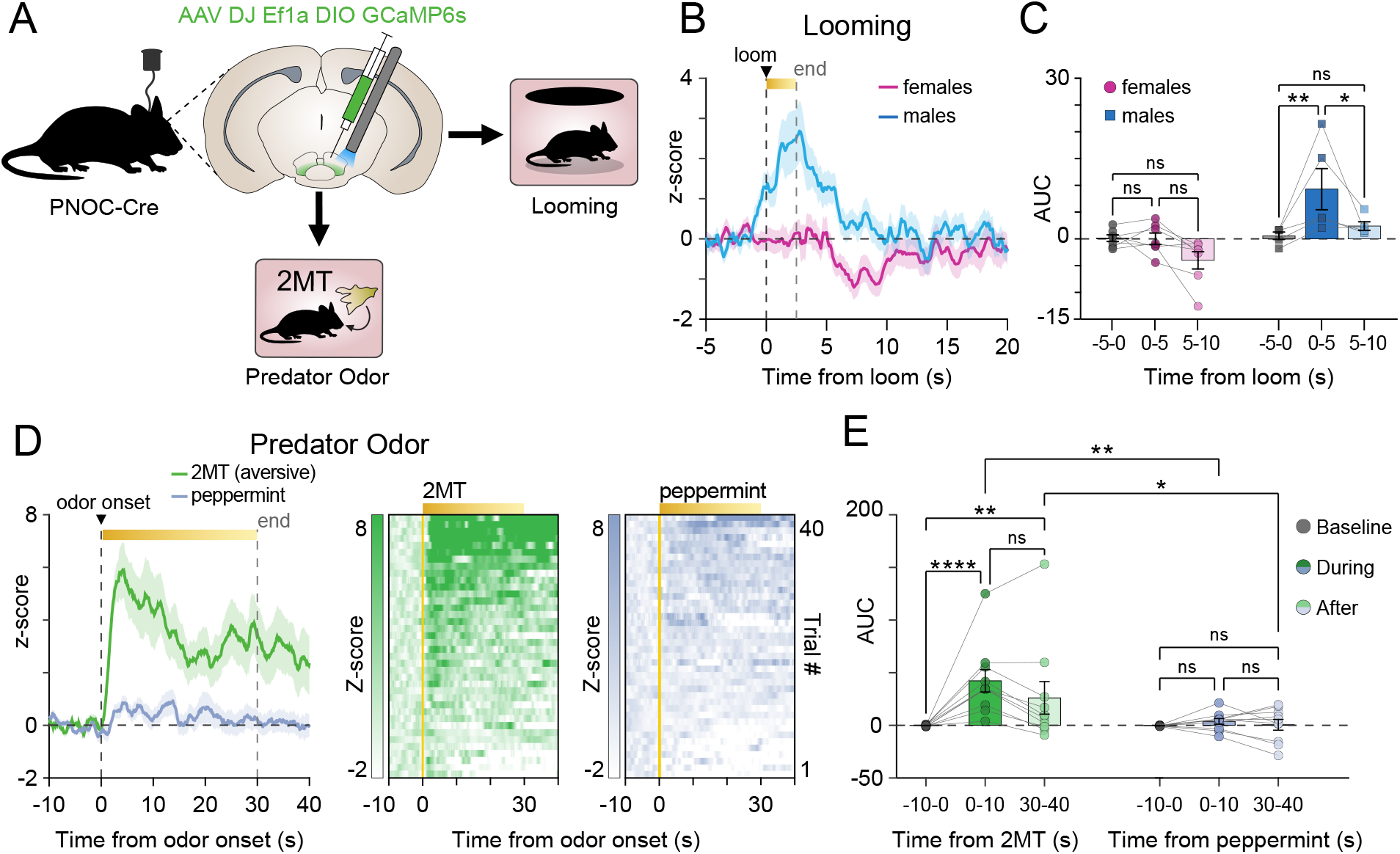
pnVTA^PNOC^ neuron activity is recruited in response to predatory threat. **A** Cartoon of DIO-GCaMP6s (GCaMP6s) injection and fiber implant into pnVTA of PNOC-Cre mice. GCaMP6s activity was recorded during looming or exposure to predator odor. **B** Averaged traces of pnVTA^PNOC^ GCaMP6s activity for males (blue, N = 5 mice) and females (magenta, N = 7 mice) aligned to looming onset. **C** Area under the curve (AUC) for averaged traces from B for females (left, magenta) and males (right, blue), calculated over 5-second intervals. GCaMP6s activity increases during and immediately after looming in males, but not females (2-way ANOVA with Tukey’s multiple comparisons test, **p < 0.01, *p < 0.05, N = 5 males, 7 females). **D** Left: Averaged traces of pnVTA^PNOC^ GCaMP6s activity surrounding 30-second exposure to either predator odor (2% 2MT, green) or a control non-predator odor (2% peppermint oil, blue). Right: Heat map of GCaMP6s fluorescence during same epoch, each row correspond to a trial in the averaged trace (left). (N = 10 mice). **E** AUC for averaged traces from D calculated over 10-second intervals surrounding exposure to either the 2MT predator odor (left, green) or the peppermint oil (right, blue). pnVTA^PNOC^ GCaMP6s activity increases during 2MT but not peppermint oil exposure (2-way ANOVA with Tukey’s multiple comparisons test, ****p < 0.0001, **p < 0.01, *p < 0.05, N = 10 mice). All data represented as mean ± SEM.

### The lateral hypothalamus provides direct GABAergic and glutamatergic input onto pnVTA^PNOC^ neurons

The VTA receives heterogeneous inputs from stress-related regions throughout the brain, so we next investigated which afferent pathways modulate this stress-sensitive pnVTA^PNOC^ subpopulation. The lateral hypothalamus (LH) sends dense and diverse projections to the VTA ^41,42^, and is broadly involved in motivation, aversion, and the stress response ^29,43,44,45,46,47^. In line with prior tracing studies ^41^, our injection of AAV2-retro-Ef1a-DIO-eYFP into the pnVTA of VGAT-Cre or VGLUT2-Cre mice effectively labeled GABAergic and glutamatergic neurons in the LH that project to the pnVTA (**Supplementary Figure 2**).

We next determined whether these LH GABA and glutamate (LH^GABA/Glut^) inputs have synaptic connectivity onto pnVTA^PNOC^ neurons (**Figure 4**). Using PNOC-Cre mice, we injected AAV5-CaMKII-ChR2-eYFP bilaterally into the LH to allow for optogenetic control of LH^GABA/Glut^ neurons and then injected AAV5-Ef1a-DIO-mCherry bilaterally into the pnVTA to label PNOC neurons for whole-cell voltage clamp recordings (**Figure 4A,B**). At 4-6 weeks post injection, mice were sacrificed and coronal brain slices were prepared for *ex vivo* electrophysiological studies.

**FIGURE 4.**
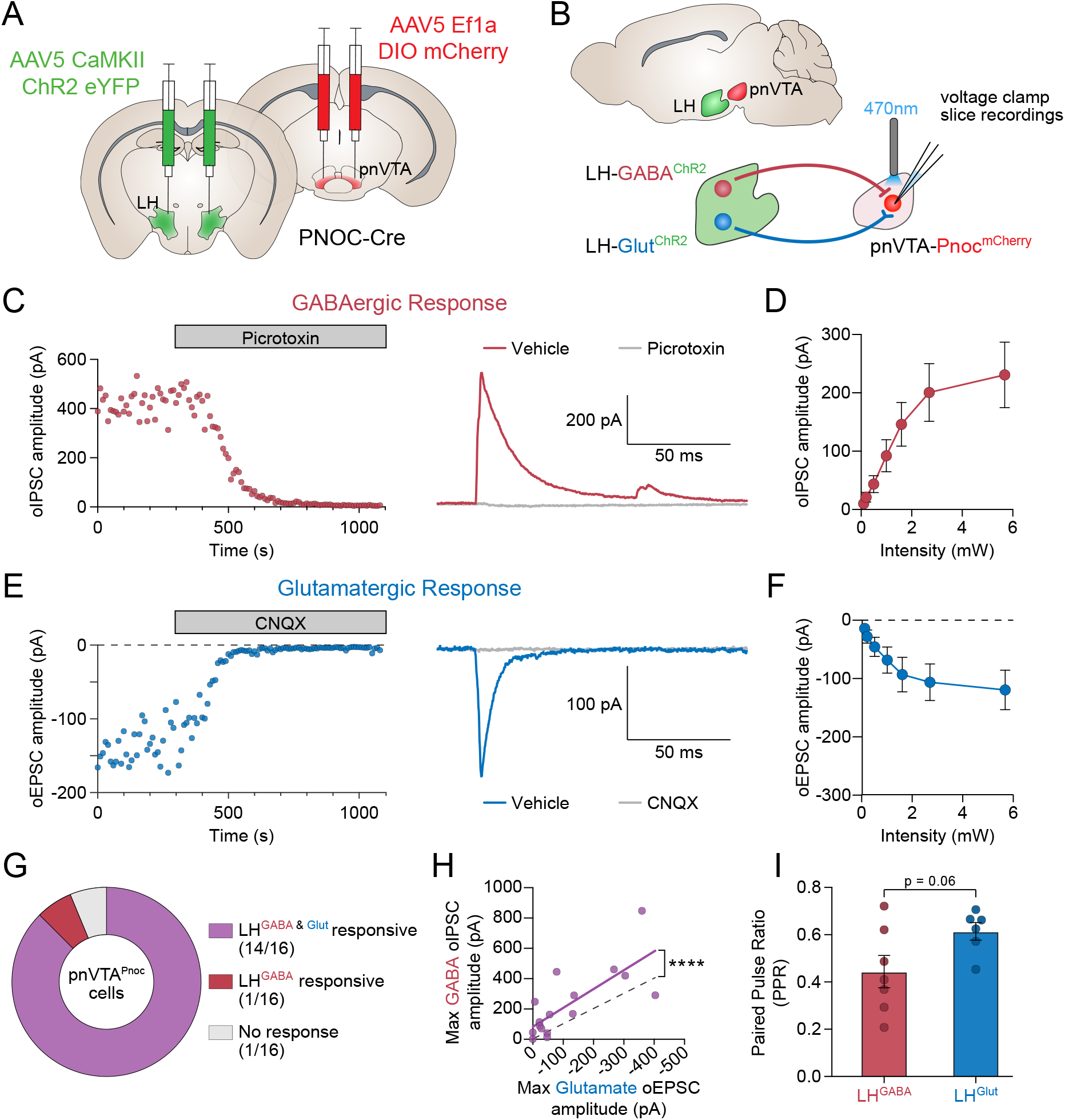
GABA and glutamatergic projections from the lateral hypothalamus provide input onto pnVTA^PNOC^ neurons. **A** Cartoon of bilateral CaMKII-ChR2-eYFP injection into lateral hypothalamus (LH) and DIO-mCherry injection into pnVTA of PNOC-Cre mice. **B** Schematic for voltage-clamp recordings of oIPSCs or oEPSCs from LH^GABA^ and LH^Glut^ terminals in the pnVTA. **C** Example trace and effect of 50 μM picrotoxin on oIPSC amplitude recorded from pnVTA^PNOC^ neurons (n = 16, N = 3). **D** Optically-evoked input/output curve of oIPSCs recorded from pnVTA^PNOC^ neurons at increasing LED stimulation intensities (n = 16, N = 3). **E** Example trace and effect of 10 μM CNQX on oEPSC amplitude recorded from pnVTA^PNOC^ neurons (n = 16, N = 3). **F** Optically-evoked input/output curve of oEPSCs recorded from pnVTA^PNOC^ neurons at increasing LED stimulation intensities (n = 16, N = 3). **G** Summary of excitatory and inhibitory responses to LH terminal stimulation from all recorded pnVTA^PNOC^ neurons. **H** Maximum oIPSC amplitude vs maximum oEPSC amplitude recorded from pnVTA^PNOC^ neurons. Nonlinear regression using a least squares fit (purple line) and null hypothesis (no GABA or glutamate amplitude bias, dashed line) are shown. (n = 16 cells per group, ****p < 0.0001). **I** Paired pulse ratio (PPR) of oIPSCs from LH^GABA^ inputs (red) and oEPSCs from LH^Glut^ inputs (blue) (n = 6-7 cells per group; two-tailed t-test, p = 0.0607).

To isolate both glutamatergic and GABAergic input from the LH within the same PNOC neurons, we used a Cesium based internal solution to record optically-evoked inhibitory and excitatory postsynaptic currents (oIPSCs and oEPSCs) at +10mV and −70mV, respectively (see Supplementary Methods). Photo-stimulation (20Hz, 2ms pulse width, 0.1–6mW) of LH terminals in the pnVTA reliably evoked oIPSCs in mCherry-labeled pnVTA^PNOC^ neurons held at +10mV that could be blocked by the GABA_A_ receptor antagonist picrotoxin (**Figure 4C, D**). Similarly, we recorded oEPSCs in pnVTA^PNOC^ neurons held at −70mV following LH terminal stimulation, which were blocked by the competitive AMPA receptor antagonist CNQX (**Figure 4E, F**). While the majority of pnVTA^PNOC^ neurons recorded responded to both LH^GABA^ and LH^Glut^ stimulation (**Figure 4G**), we found that the magnitude of the inhibitory response was significantly greater than the excitatory response (**Figure 4H**). This was accompanied by a trend toward a lower paired pulse ratio (PPR) for the inhibitory responses, suggesting higher release probability and stronger synaptic input (**Figure 4I**).Taken together, these data suggest that the lateral hypothalamus, a key nucleus in orchestrating stress-related behaviors, provides robust GABAergic and glutamatergic input to pnVTA^PNOC^ neurons, suggesting that these pathways could influence the VTA’s responsivity to stress.

After identifying synaptic connectivity between the LH and pnVTA^PNOC^ neurons, we investigated the functional activity of these LH terminals in the pnVTA during stress exposure first by recording LH^GABA^ calcium activity during exposure to predator odor (**Supplementary Figure 3A, B**). While our earlier findings showed an increase in pnVTA^PNOC^ activity in response to predator odor, when recording from LH^GABA^ terminals we instead observed a decrease in calcium activity (**Supplementary Figure 3C,D**, 1-way ANOVA, F_4,8_=4.593, p=0.0321). To further evaluate LH function in the pnVTA during stress, we used an optogenetic approach to stimulate LH^Glut^ terminal activity during the tail suspension test (TST) ^35^. We observed a general trend toward less time spent struggling during the TST when mice received continuous 20Hz stimulation of LH^Glut^ terminals compared to control days, indicative of increased anhedonia (**Supplementary Figure 3G**). Together, these results indicate a functional role for upstream LH projections to the pnVTA as a mechanism for responding to stress.

## DISCUSSION

Our findings reveal new insights into a pathway by which stress interfaces with a neuropeptide subpopulation known to direct motivated reward-seeking behaviors through its influence on mesolimbic circuitry. We demonstrate that pnVTA^PNOC^ neurons are selectively activated in response to stress rather than salience in a primarily non-sex-dependent manner, except during looming predator stress where activity increased in males only. Our findings also indicate that pnVTA^PNOC^ neurons have varying sensitivity to different stressors, with physical and predator-based stimuli producing a larger dynamic response than the innately stressful environmental cues experienced during exploratory behaviors. Whether these differences in dynamics are related to the perceived valence of the stressful stimulus will be an important follow-up for further study.

N/OFQ signaling has been implicated in the stress response, but the observed effects of stress on the N/OFQ-NOPR system vary widely both across the brain and depending on the form and duration of stress exposure ^28,48^. Prior studies have also reported notable sex differences in rats, where stress-induced changes to N/OFQ signaling were more prominent in males ^49,50^. While most of the stressors we evaluated in this study evoked similar pnVTA^PNOC^ responses in males and females, the activity during looming that occurred exclusively in male mice mirrors these prior findings where a stress-evoked change in N/OFQ signaling was only seen in males. On the contrary, we observed a potentiation of pnVTA^PNOC^ activation during foot shock in females relative to males (**Supplementary Figure 1**), suggesting sex-dependent effects in N/OFQ’s response to stress can also extend to females. Beyond sex-based differences, it would be interesting to determine whether individual heterogeneity in terms of an animal’s resilience or vulnerability to stress further shapes how pnVTA^PNOC^ neurons respond during stress exposure.

The N/OFQ-NOPR system has emerged as a promising therapeutic target for anhedonia. Preclinical studies with NOPR antagonists have repeatedly shown them to exhibit an antidepressant-like profile, and one antagonist evaluated in a clinical trial was found to improve depression scores, although it did not reach the predefined efficacy criteria (required >88% efficacy relative to placebo; achieved 82.9%) ^51^. Despite the large body of evidence demonstrating that NOPR inhibition induces an antidepressant-like phenotype (for a summary see Gavioli and Calo, 2013), the specific circuit-level mechanisms by which blockade of N/OFQ signaling improves anhedonic symptoms are less understood. Previous work reported that inhibition of pnVTA^PNOC^ activity enhances effort-based reward seeking in mice ^25^, suggesting a mechanism by which N/OFQ signaling may regulate anhedonic behaviors. It is well established that stress is a prominent factor in the pathophysiology and development of anhedonia, and our findings in this study identify a pathway through which stress could suppress motivation by driving excessive activation of N/OFQ signaling in the pnVTA, which could be then reversed by NOPR antagonism. Further work with specialized genetic approaches is needed to more directly test this hypothesis however, particularly in evaluating whether inhibition of pnVTA^PNOC^ activity can improve chronic stress-induced motivation deficits.

Clinical studies have found increased plasma levels of N/OFQ in patients with major depressive disorder (MDD), post-partum depression, and bipolar depression, all of which are commonly associated with anhedonic behaviors and reduced motivation ^52,53^. Interestingly, plasma N/OFQ levels are suppressed relative to healthy controls in patients with bipolar mania ^53^, which is more often associated with increased impulsivity and higher reward-seeking drive. This suggests that N/OFQ signaling may be critical for fine-tuning reward-related behavior, and imbalances in either direction of standard expression levels could drive opposing effects on motivation. Our prior findings also support this idea, as we saw that over-activation of pnVTA^PNOC^ neurons suppressed reward-seeking behavior while pnVTA^PNOC^ inhibition enhanced motivation ^25^. Our findings here showed pnVTA^PNOC^ activation in response to stress, thus we would predict that stress-induced activation of this population would contribute to an overall anhedonic phenotype. However, it is important to note that our findings are still limited by the selection of stressors used in this study. Given that stress exposure does not always induce an anhedonic phenotype and can instead exacerbate impulsive behaviors such as drug self-administration ^54,55^, further work should investigate if other forms and extended durations of stress have a suppressive effect on pnVTA^PNOC^ activity.

The upstream circuitry that engages the pnVTA^PNOC^ subpopulation was unknown. The lateral hypothalamus (LH) is a critical hub with roles in stress and reward-seeking that innervates the pnVTA with both excitatory and inhibitory inputs ^41^, making it promising candidate for pnVTA^PNOC^ afferent modulation. In addition to the LH, our retrograde tracing study identified several additional anatomical targets with known roles in responding to or coping with stress that provide input to the pnVTA, such as the ventrolateral periaqueductal gray ^56^, lateral habenula ^57^, and nucleus accumbens shell ^58^. While we chose to focus on the LH in this study, the functional connectivity of these other notable regions onto pnVTA^PNOC^ neurons during stress responses warrants further exploration.

Our patch-clamp recordings identify synaptic connectivity from both GABAergic and glutamatergic inputs onto pnVTA^PNOC^ neurons (**Figure 4, Supplementary Figure 2**) and indicate a functional role of LH glutamatergic activity at the pnVTA in driving anhedonic behavior (**Supplementary Figure 3**). These findings align with other investigations of LH projections to the VTA, where LH glutamatergic activity has been shown to drive avoidance behaviors and LH GABAergic activity to promote positive reinforcement ^29,46^. Our recordings of LH^GABA^ presynaptic terminal activity during predator odor stress indicate that suppression of LH^GABA^ activity could serve as a disinhibitory mechanism that allows for heightened activation of pnVTA^PNOC^ neurons in response to stress, however thus far we have only observed a trend toward LH^GABA^ suppression.

It is also important to note that the electrophysiological studies used a CaMKII-driven channelrhodopsin to target both GABA and glutamate LH populations simultaneously, and separation of oEPSCs and oIPSCs was achieved through a voltage-clamp approach. Additional experiments could isolate the synaptic input from GABAergic and glutamatergic LH inputs separately, and/or use Cre and FLP-based transgenic methods to more selectively target other unique sub-types of LH neurons. Furthermore, given the prior evidence of LH peptidergic neuromodulation of motivated behaviors within the VTA, future studies should evaluate the effect of LH neuropeptide release, particularly the peptides orexin, hypocretin, and neurotensin ^30,59,60,61^, on pnVTA^PNOC^ activity to identify if there are multi-transmitter effects that can recruit or dampen these neurons.

In conclusion, the findings we report here provide insights into how stress modulates N/OFQ signaling in the VTA and reveal the involvement of LH inputs in driving these responses. These results advance our understanding of how the N/OFQ system interfaces with both stress and motivational deficits within a single circuit, laying the groundwork for further studies to explore stress-related mechanisms for anhedonic behavior in the context of this neuropeptidergic pathway. Future research should examine the long-term effects of chronic stress on pnVTA^PNOC^ activity and evaluate the therapeutic potential of NOPR antagonism within this circuit during motivation in models of stress-induced anhedonia.

## Abbreviations

Pnoc: prepronociceptin
N/OFQ: nociceptin/orphanin FQ
VTA: ventral tegmental area

## ACKNOWLEDGMENTS

We would like to thank Drs. Azra Suko, Dr. Selena Schrieber and Larry Zweifel for their help with virus production, and Dr. Raaj Gowrishankar for assistance with fiber photometry analysis and interpretation.

## AUTHOR CONTRIBUTIONS

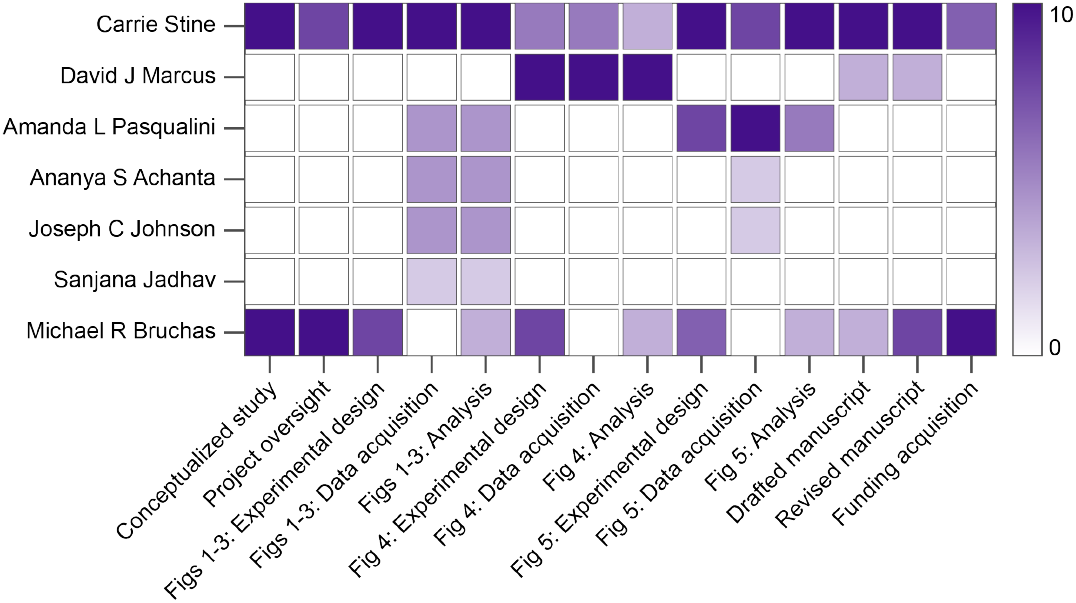

## DATA AVAILABILITY

The datasets generated and analyzed in this study are available from the corresponding author upon reasonable request.

## SUPPORTING INFORMATION

Additional supporting information may be found in the online version of the article.

## SUPPLEMENTARY METHODS

### Fiber photometry

531-Hz sinusoidal LED light (Thorlabs, LED light: M470F3; LED driver: DC4104) was bandpass filtered (470 ± 20nm, Doric, FMC4) to excite GCaMP6s, a 211-Hz sinusoidal LED light (Thorlabs, M405FP1; LED driver: DC4104) was bandpass filtered (405 ± 10nm, Doric, FMC4) to evoke Ca^2+^-independent isosbestic control emission. LED intensities were measured at the tip of the optic fiber and adjusted to 30µW before each recording. GCaMP6s emission traveled back through the same optic fiber then was bandpass filtered, (525 ± 25nm, Doric, FMC4), detected by a photoreceiver (Doric, DFD_FOA_FC), and recorded by a real-time processor (TDT, RZ5P). For the ChrimsonR stimulation experiments, a 635nm laser was passed through the filter cube at 1mW intensity to deliver red light at the tip of the optic fiber ^32^.

### Fiber photometry analysis

Custom MATLAB scripts were used to detrend decay from bleaching by fitting a 4^th^ degree polynomial function to the raw signal (470nm) and isosbestic (405nm) traces, then dividing by the resulting curve. Then signal was normalized by dividing an LLS fit of the isosbestic trace scaled to the signal. The processed traces were then down-sampled by a factor of 100, smoothed across a rolling 1s window, extracted in windows surrounding the onset of relevant behavioral events (tail lift, odor, shock, air puff, looming, open arm entry, center entry), z-scored relative to the mean and standard deviation of each event window, and then averaged.

### Electrophysiology

Coronal brain slices were prepared at 250 μM on a vibrating Leica VT1000S microtome using standard procedures. Mice were anesthetized with Isoflurane, and transcardially perfused with an ice-cold, oxygenated cutting solution consisting of (in mM): 93 N-Methyl-D-glucamine (NMDG), 2.5 KCL, 20 HEPES, 30 NaHCO3, NaH2PO4, 10 MgSO4⍰7H20, 0.5 CaCl2⍰2H20, 25 glucose, 3 Na^+^-pyruvate, 5 Na^+^-ascorbate, and 5 N-acetylcysteine. Following collection of coronal sections, the brain slices were transferred to a 34°C chamber containing oxygenated cutting solution for a 10-minute recovery period. Slices were then transferred to a holding chamber consisting of (in mM) 92 NaCl, 2.5 KCl, 20 HEPES, 2 MgSO4⍰7H20, 1.2 NaH2PO4, 30NaHCO3, 2 CaCl2⍰2H20, 25 glucose, 3 Na-pyruvate, 5 Na-ascorbate, 5 N-acetylcysteine and were allowed to recover for ≥ 30 min. For recording, slices were perfused with oxygenated artificial cerebrospinal fluid (ACSF; 31-33°C; 300-303 milliosmoles) consisting of (in mM): 113 NaCl, 2.5 KCl, 1.2 MgSO4⍰7H20, 2.5 CaCl2⍰6H20, 1 NaH2PO4, 26 NaHCO3, 20 glucose, 3 Na^+^-pyruvate, 1 Na^+^-ascorbate, at a flow rate of 2-3ml/min. For recordings of inhibitory currents, 10 μM CNQX was added to the external solution.

mCherry-labeled pnVTA^Pnoc^ neurons were initially voltage clamped in whole-cell configuration using borosilicate glass pipettes (2-4MΩ). For recordings of excitatory currents, pipettes were filled with internal solution containing (in mM): 125 K^+^-gluconate, 4 NaCl, 10 HEPES, 4 MgATP, 0.3 Na-GTP, and 10 Na-phosphocreatine (pH 7.30-7.35). The patch pipette also included 50 μM picrotoxin to block GABA_A_ currents. For recordings of inhibitory currents, pipettes were filled with 115 CsCl2, 5 NaCl, 10 HEPES, 5 QX-314, 4 Mg-ATP, 0.3 Na-GTP, and 10 Glucose (pH 7.30-7.35). Following break-in to the cell, we waited ≥ 3 minutes to allow for exchange of internal solution and stabilization of membrane properties. Neurons with an access resistance of > 30MΩ or that exhibited greater than a 20% change in access resistance during the recording were not included in our datasets. For all voltage clamp experiments, neurons were held at −70mV.

### Ex vivo optogenetics

PNOC-Cre mice were injected with 300 nL of AAV5-CaMKII-ChR2(H134R)-eYFP bilaterally into the LH, and 300 nL of AAV5-EF1a-DIO-mCherry was injected bilaterally into the pnVTA to allow us to record from PNOC^+^ neurons in the pnVTA. 3-5 weeks of viral expression was allowed prior to sacrificing the mice. For optogenetic recordings of input/output curves, we used a Thorlabs LEDD1B T-Cube driver and obtained separate recordings of 470nm wavelength oEPSCs / oIPSCs at 7 output levels corresponding to 5.7, 2.7, 1.6, 1, 0.5, 0.2, and 0.1 mW of LED intensity. Paired pulse ratio (PPR) recordings of oEPSCs / oIPSCs were obtained in voltage-clamp with an inter-stimulus interval of 50ms. PPR is reported as a ratio between the amplitude of the second oEPSC / oIPSC divided by the first. A light exposure time of 2ms was used for all optogenetic experiments.

### Immunohistochemistry

Free-floating brain sections were washed in 0.1 M PBS for 3 x 10 min intervals. Sections were then placed in blocking buffer (0.5% Triton X-100 and 5% natural goat serum in 0.1 M PBS) for 1 hour at room temperature. After blocking buffer, sections were placed in primary antibody (chicken anti-GFP, 1:2000, Abcam) overnight at 4°C. After 3 x 10 min 0.1 M PBS washes, sections were incubated in secondary antibody (AlexaFluor 488 goat anti-chicken, Abcam) for 2 hours at room temperature, followed by another round of washes (3 x 10 min in 0.1 M PBS, 3 x 10 min in 0.1 M PBS). After immunostaining, sections were mounted and coverslipped with Vectashield HardSet mounting medium containing DAPI (Vector Laboratories) and imaged on a Leica DM6 B microscope.

## SUPPLEMENTARY FIGURES

**SUPPLEMENTARY FIGURE 1.**
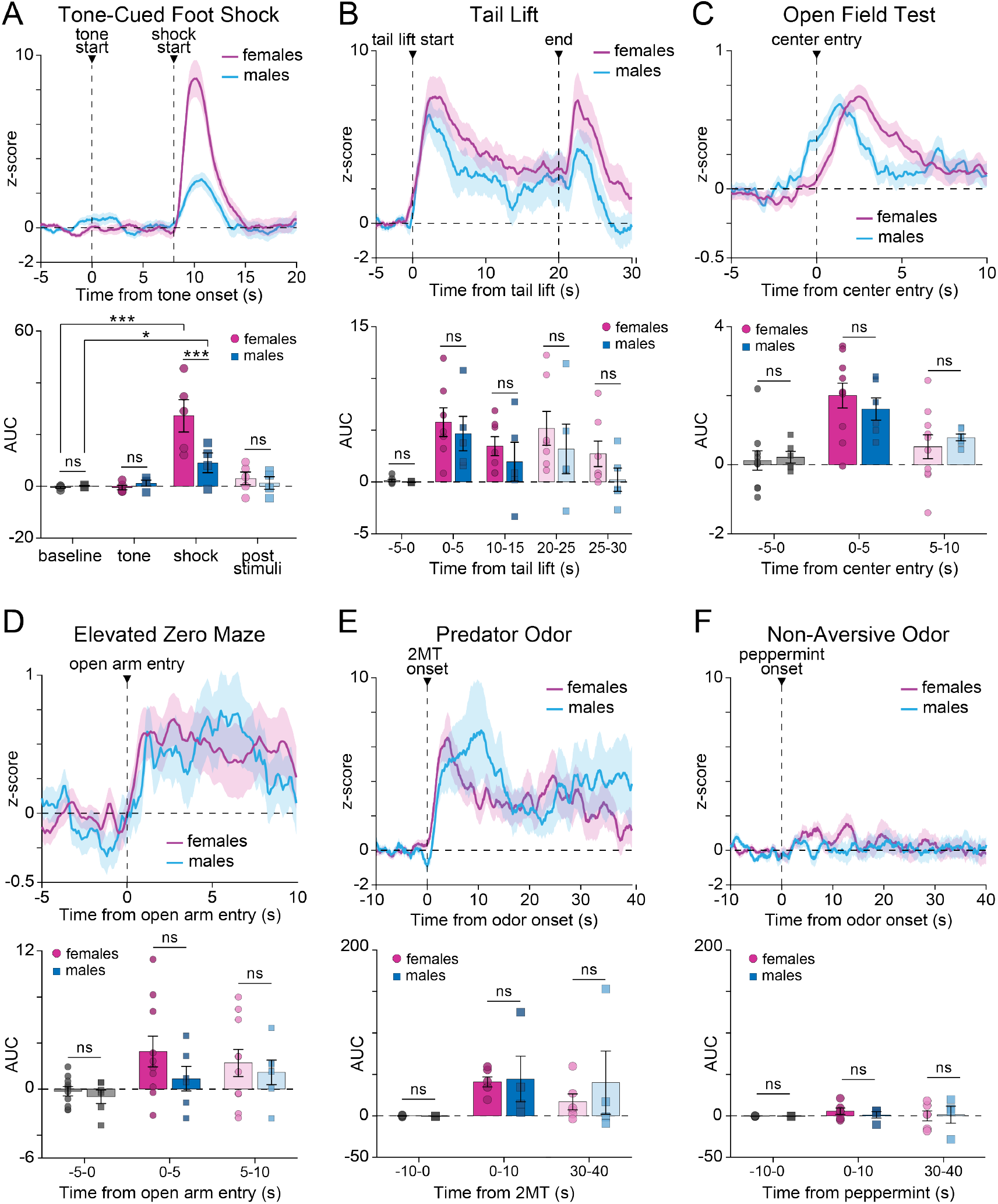
Stress engages pnVTA^PNOC^ neurons in both male and female mice. **A** Top: Averaged trace of pnVTA^PNOC^ GCaMP6s activity during epoch surrounding tone-cued foot shock, aligned to tone onset in male (blue) and female (magenta) mice. Bottom: Area under the curve (AUC) for averaged traces, calculated over 8-second intervals surrounding cued-foot shock events for males (blue) and females (magenta). GCaMP6s signal increases in response to shock but not tone for both sexes (1-way ANOVA with Tukey’s multiple comparison test, ***p < 0.001 *p < 0.05, N = 4 males, 5 females), although the magnitude of the increase is larger in females (2-way ANOVA with Tukey’s multiple comparisons test, ***p < 0.001, N = 4 males, 5 females). **B–F** Same as A for **B** 20s tail lift (2-way ANOVA p=0.3767, N = 5 males, 7 females), **C** open field test center entry (2-way ANOVA p=0.9794, N = 6 males, 10 females), **D** elevated zero maze open arm entry (2-way ANOVA p=0.3212, N = 6 males, 10 females), **E** predator (2% 2MT) odor (2-way ANOVA p=0.7345, N = 4 males, 6 females), and **F** non-aversive (2% peppermint oil) odor (2-way ANOVA p=0.7633, N = 4 males, 6 females). All data represented as mean ± SEM.

**SUPPLEMENTARY FIGURE 2.**
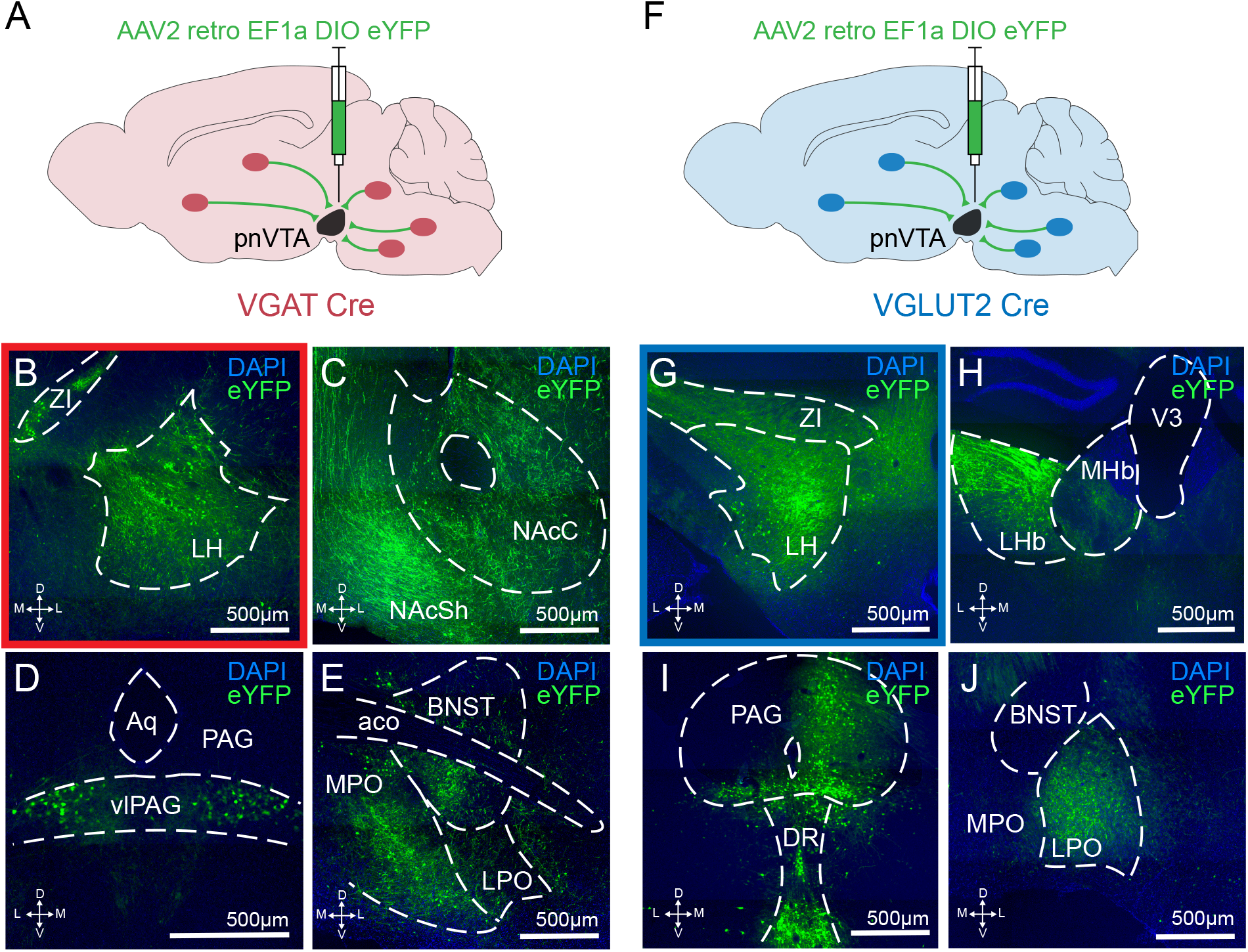
Retrograde mapping of GABA and glutamatergic inputs to the pnVTA. **A** Cartoon showing injection of a retrograde DIO-eYFP into the pnVTA of VGAT-Cre mice (N = 2). Slices across the entire brain were imaged to identify sources of GABAergic input to the pnVTA. **B–E** Representative images from brain regions with relatively high expression of retrograde eYFP (eYFP, green; DAPI, blue). **F–J** Same as A–E but in VGLUT2-Cre mice (N = 2) to identify sources of glutamatergic input to the pnVTA. LH = lateral hypothalamus, ZI = zona incerta, NAcC = nucleus accumbens core, NAcSh = nucleus accumbens shell, (vl)PAG = (ventro-lateral) periaqueductal gray, Aq = cerebral aqueduct, BNST = bed nuclei of the stria terminalis, MPO = medial preoptic area, LPO = lateral preoptic area, aco = anterior commissure, LHb = lateral habenula, MHb = medial habenula, V3 = third ventricle, DR = dorsal raphe.

**SUPPLEMENTARY FIGURE 3.**
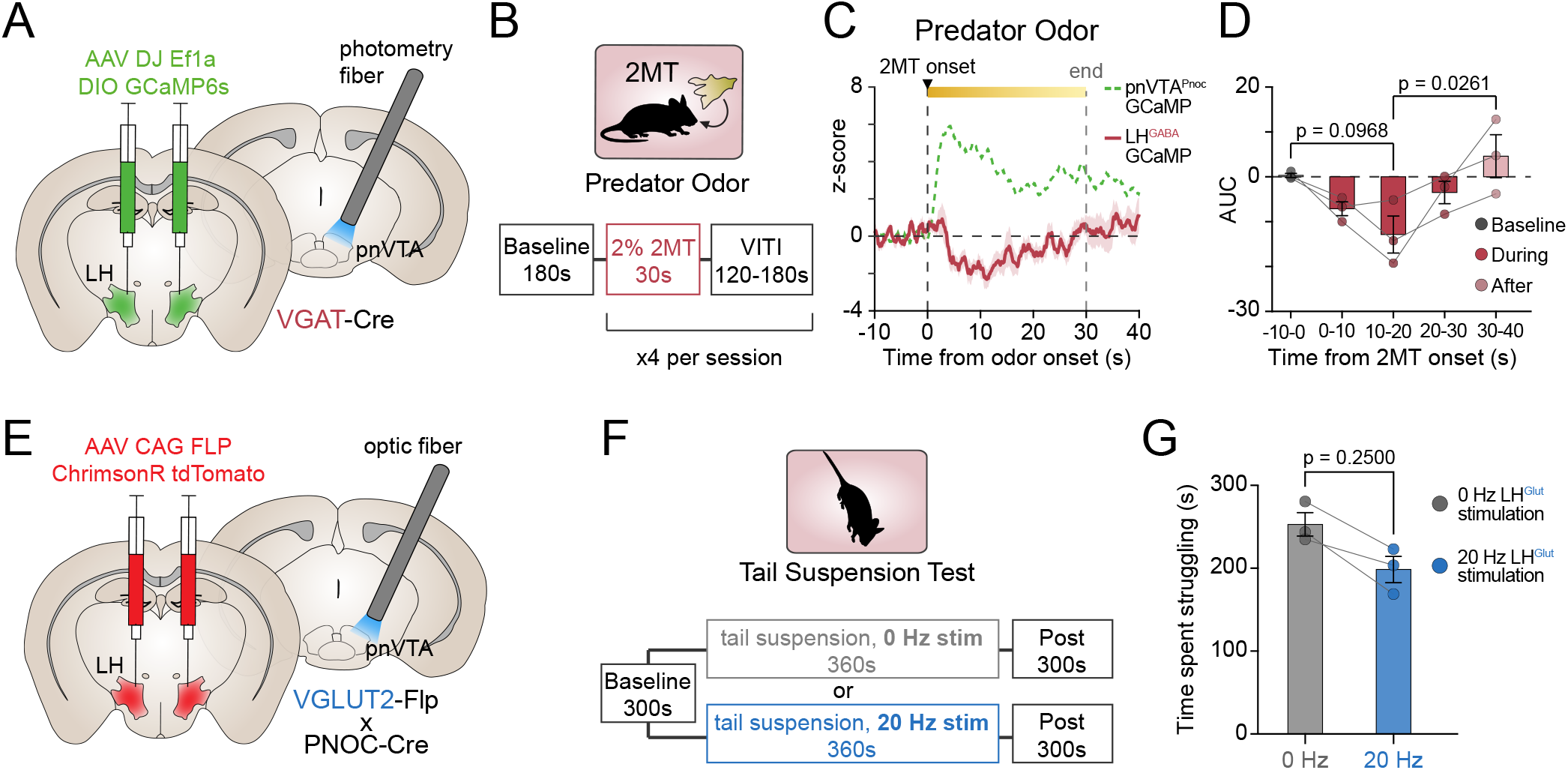
Inhibitory lateral hypothalamic input to the pnVTA disengages during stress exposure. **A** Cartoon of bilateral DIO-GCaMP6s injection into lateral hypothalamus (LH) with fiber implant above the pnVTA in VGAT-Cre mice. **B** LH^GABA^ terminals in the pnVTA were recorded during exposure to predator odor (2% 2MT, same trial structure as in Figure 2). **C** Averaged trace of LH^GABA^ terminal activity in the pnVTA during predator odor exposure (red). Averaged pnVTA^PNOC^ GCaMP6s trace from Figure 2 is shown in the green dotted line for comparison. **D** Area under the curve (AUC) for averaged traces from C for pnVTA^PNOC^ neurons (green) and LH^GABA^ terminals in the pnVTA (red). LH^GABA^ terminals are suppressed during 2MT exposure while pnVTA^PNOC^ neurons become activated (2-way ANOVA with Tukey’s multiple comparisons test, **p < 0.01, N = 10 mice, pnVTA^PNOC^; N = 3 mice, LH^GABA^). **E** Cartoon of bilateral FLP-ChrimsonR-tdTomato injection in the LH with fiber implant above the pnVTA in VGLUT2-Flp x PNOC-Cre mice. **F** Trial structure for the tail suspension test (TST). Mice were suspended for 6 minutes and received either 0 or 20 Hz ChrimsonR stimulation (1mW power, cycling ON for 5s and OFF for 15s throughout the 6-minute suspension) in counterbalanced sessions. **G** Total time spent struggling during the TST for mice receiving 0 Hz (gray) or 20 Hz (blue) stimulation of LH^Glut^ terminals in the pnVTA (two-tailed Wilcoxon test, p = 0.25, N = 3 mice). All data represented as mean ± SEM.

## SUPPLEMENTARY TABLES

**SUPPLEMENTARY TABLE 1.**
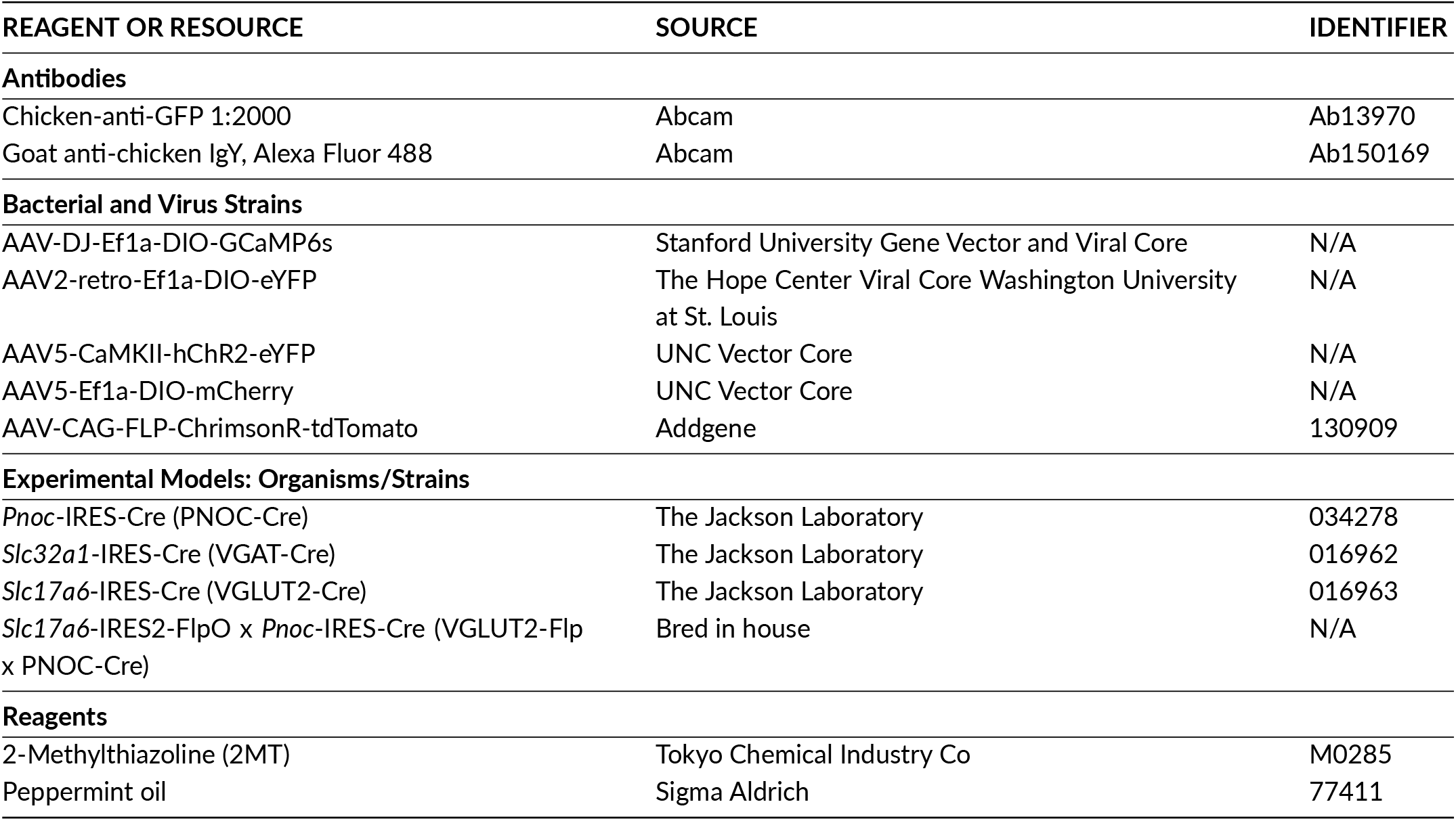
Key resources.

**SUPPLEMENTARY TABLE 2.**
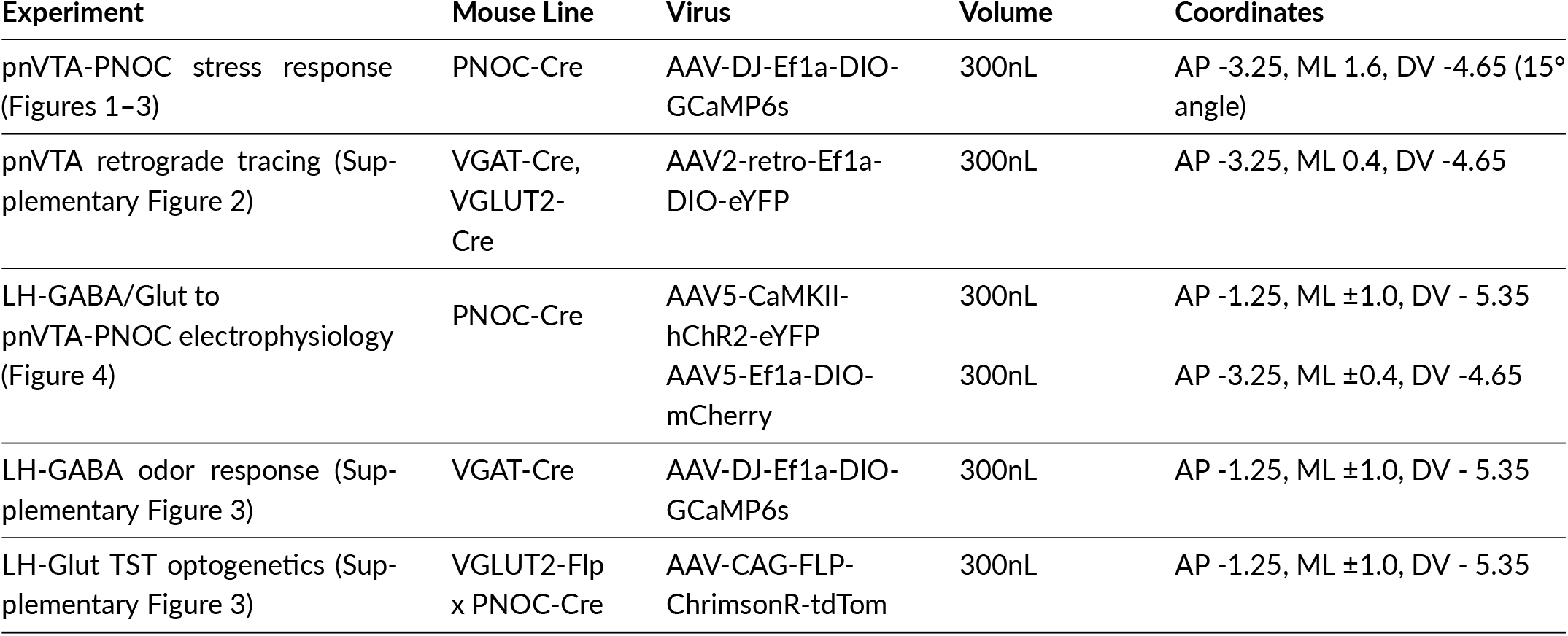
Viruses, injection coordinates, and implant types.

| Experiment | Mouse Line | Virus | Volume | Coordinates |
| --- | --- | --- | --- | --- |
| pnVTA-PNOC stress response (Figures 1–3) | PNOC-Cre | AAV-DJ-Ef1a-DIO-GCaMP6s | 300nL | AP -3.25, ML 1.6, DV -4.65 (15° angle) |
| pnVTA retrograde tracing (Supplementary Figure 2) | VGAT-Cre, VGLUT2-Cre | AAV2-retro-Ef1a-DIO-eYFP | 300nL | AP -3.25, ML 0.4, DV -4.65 |
| LH-GABA/Glut to pnVTA-PNOC electrophysiology (Figure 4) | PNOC-Cre | AAV5-CaMKII-hChR2-eYFP | 300nL | AP -1.25, ML $\pm 1.0$, DV - 5.35 |
| | | AAV5-Ef1a-DIO-mCherry | 300nL | AP -3.25, ML $\pm 0.4$, DV -4.65 |
| LH-GABA odor response (Supplementary Figure 3) | VGAT-Cre | AAV-DJ-Ef1a-DIO-GCaMP6s | 300nL | AP -1.25, ML $\pm 1.0$, DV - 5.35 |
| LH-Glut TST optogenetics (Supplementary Figure 3) | VGLUT2-Flp x PNOC-Cre | AAV-CAG-FLP-ChrimsonR-tdTom | 300nL | AP -1.25, ML $\pm 1.0$, DV - 5.35 |

