## Supplementary Information for "Identification of a stress-sensitive endogenous opioid-containing neuronal population in the paranigral ventral tegmental area"

### Supplementary Figures

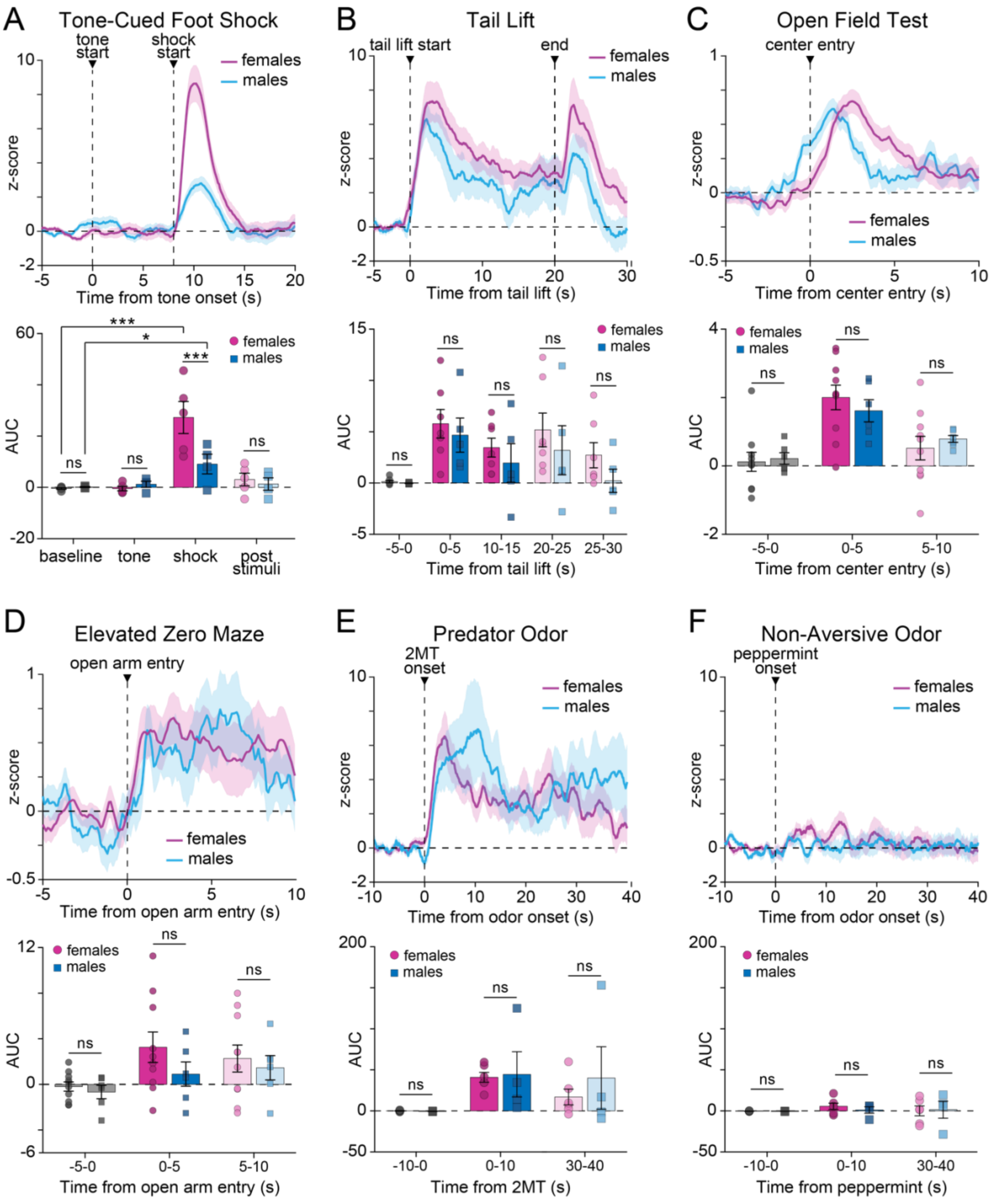

**Supplementary Figure 1: Stress engages pnVTA$^{PNOC}$ neurons in both male and female mice.**

**A** Top: Averaged trace of pnVTA$^{PNOC}$ GCaMP6s activity during epoch surrounding tone-cued foot shock, aligned to tone onset in male (blue) and female (magenta) mice. Bottom: Area under the curve (AUC) for averaged traces, calculated over 8-second intervals surrounding cued-foot shock events for males (blue) and females (magenta). GCaMP6s signal increases in response to shock but not tone for both sexes (1-way ANOVA with Tukey's multiple comparison test, ***p < 0.001 *p < 0.05, N = 4 males, 5 females), although the magnitude of the increase is larger in females (2-way ANOVA with Tukey's multiple comparisons test, ***p < 0.001, N = 4 males, 5 females). **B–F** Same as A for **B** 20s tail lift (2-way ANOVA p=0.3767, N = 5 males, 7 females), **C** open field test center entry (2-way ANOVA p=0.9794, N = 6 males, 10 females), **D** elevated zero maze open arm entry (2-way ANOVA p=0.3212, N = 6 males, 10 females), **E** predator (2% 2MT) odor (2-way ANOVA p=0.7345, N = 4 males, 6 females), and **F** non-aversive (2% peppermint oil) odor (2-way ANOVA p=0.7633, N = 4 males, 6 females). All data represented as mean ± SEM.

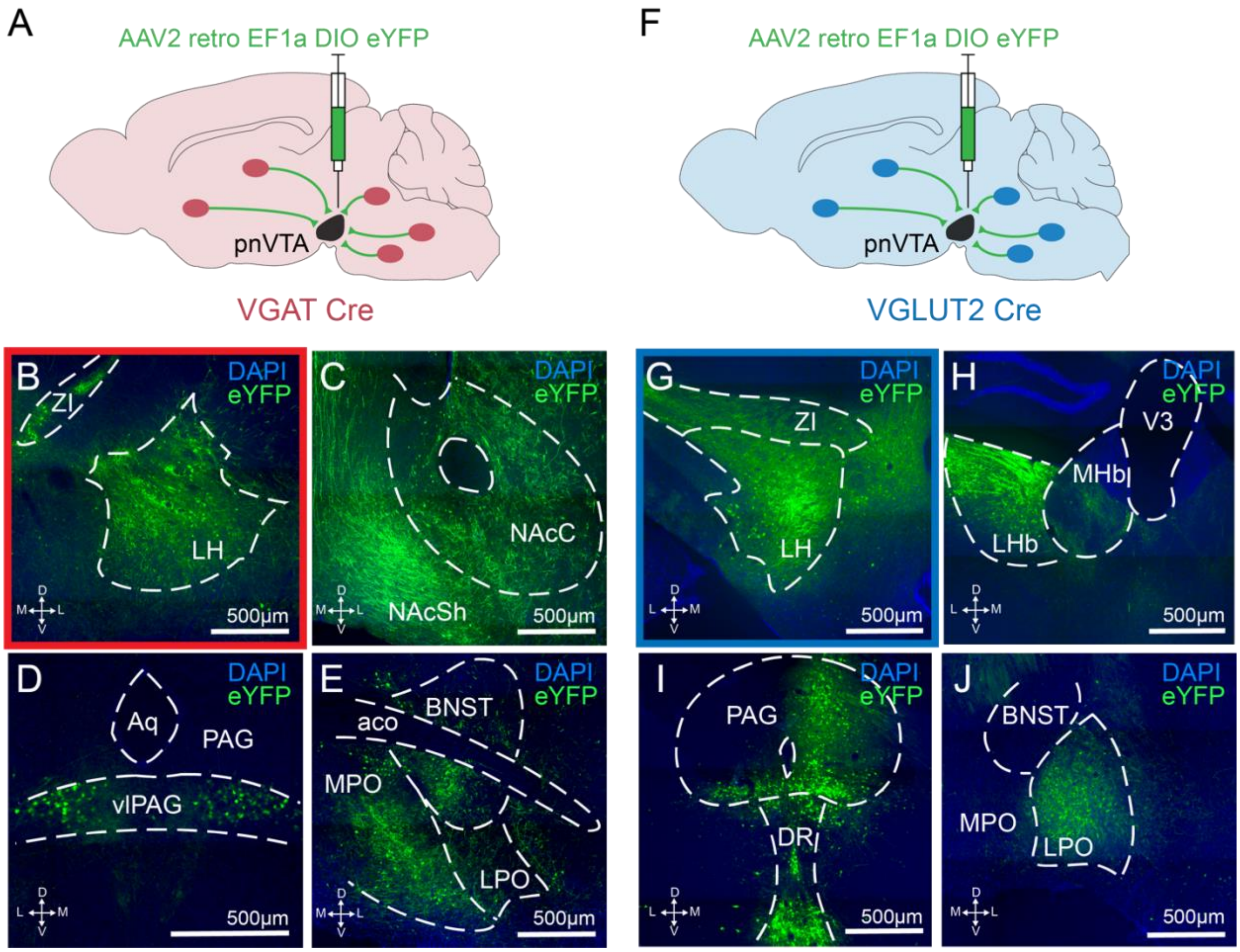


**Supplementary Figure 2: Retrograde mapping of GABA and glutamatergic inputs to the pnVTA.**

**A** Cartoon showing injection of a retrograde DIO-eYFP into the pnVTA of VGAT-Cre mice (N = 2). Slices across the entire brain were imaged to identify sources of GABAergic input to the pnVTA. **B–E** Representative images from brain regions with relatively high expression of retrograde eYFP (eYFP, green; DAPI, blue). **F–J** Same as A–E but in VGLUT2-Cre mice (N = 2) to identify sources of glutamatergic input to the pnVTA. LH = lateral hypothalamus, ZI = zona incerta, NAcC = nucleus accumbens core, NAcSh = nucleus accumbens shell, (vl)PAG = (ventro-lateral) periaqueductal gray, Aq = cerebral aqueduct, BNST = bed nuclei of the stria terminalis, MPO = medial preoptic area, LPO = lateral preoptic area, aco = anterior commissure, LHb = lateral habenula, MHb = medial habenula, V3 = third ventricle, DR = dorsal raphe.

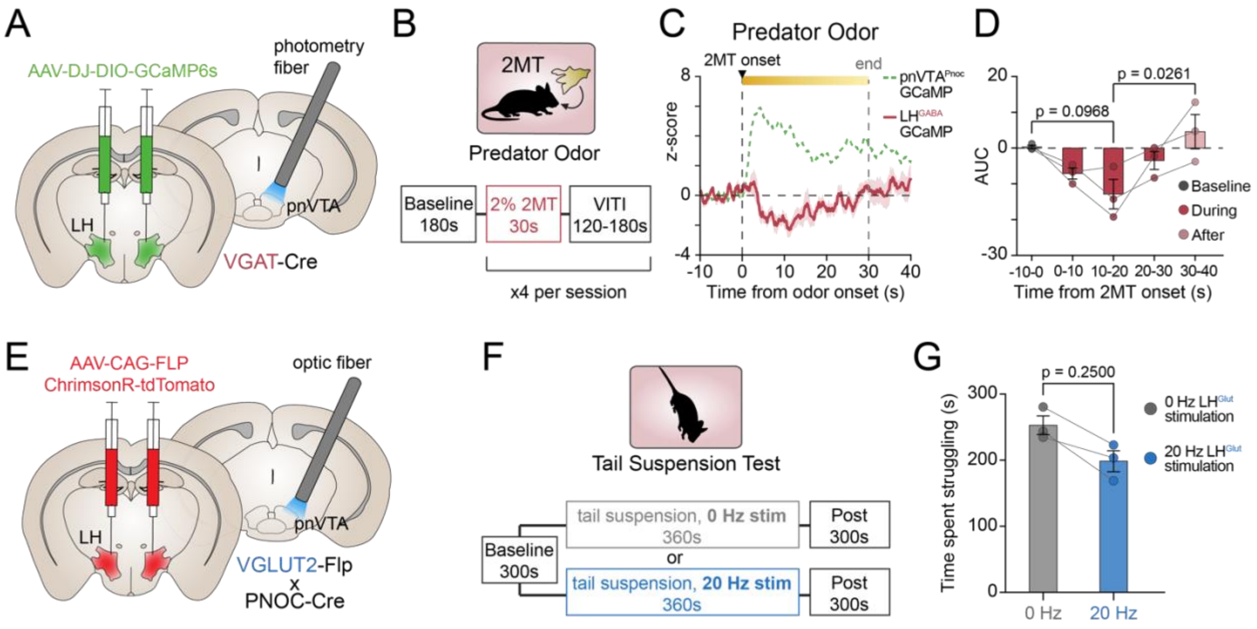


**Supplementary Figure 3: Inhibitory lateral hypothalamic input to the pnVTA disengages during stress exposure.**
**A** Cartoon of bilateral DIO-GCaMP6s injection into lateral hypothalamus (LH) with fiber implant above the pnVTA in VGAT-Cre mice. **B** $LH^{GABA}$ terminals in the pnVTA were recorded during exposure to predator odor (2% 2MT, same trial structure as in Figure 2). **C** Averaged trace of $LH^{GABA}$ terminal activity in the pnVTA during predator odor exposure (red). Averaged $pnVTA^{PNOC}$ GCaMP6s trace from Figure 2 is shown in the green dotted line for comparison. **D** Area under the curve (AUC) for averaged traces from C for $pnVTA^{PNOC}$ neurons (green) and $LH^{GABA}$ terminals in the pnVTA (red). $LH^{GABA}$ terminals are suppressed during 2MT exposure while $pnVTA^{PNOC}$ neurons become activated (2-way ANOVA with Tukey's multiple comparisons test, $^{**}p < 0.01$, N = 10 mice, $pnVTA^{PNOC}$; N = 3 mice, $LH^{GABA}$). **E** Cartoon of bilateral FLP-ChrimsonR-tdTomato injection in the LH with fiber implant above the pnVTA in VGLUT2-Flp x PNOC-Cre mice. **F** Trial structure for the tail suspension test (TST). Mice were suspended for 6 minutes and received either 0 or 20 Hz ChrimsonR stimulation (1mW power, cycling ON for 5s and OFF for 15s throughout the 6-minute suspension) in counterbalanced sessions. **G** Total time spent struggling during the TST for mice receiving 0 Hz (gray) or 20 Hz (blue) stimulation of $LH^{Glut}$ terminals in the pnVTA (two-tailed Wilcoxon test, $p = 0.25$, N = 3 mice). All data represented as mean ± SEM.

### Table S1: Key Resources

| REAGENT OR RESOURCE | SOURCE | IDENTIFIER |
|---|---|---|
| **Antibodies** | | |
| Chicken-anti-GFP 1:2000 | Abcam | Ab13970 |
| Goat anti-chicken IgY, Alexa Fluor 488 | Abcam | Ab150169 |
| **Bacterial and Virus Strains** | | |
| AAV-DJ-Ef1a-DIO-GCaMP6s | Stanford University<br>Gene Vector and Viral Core | N/A |
| AAV2-retro-Ef1a-DIO-eYFP | The Hope Center Viral Core<br>Washington University at St. Louis | N/A |
| AAV5-CaMKII-hChR2-eYFP | UNC Vector Core | N/A |
| AAV5-Ef1a-DIO-mCherry | UNC Vector Core | N/A |
| AAV-CAG-FLP-ChrimsonR-tdTomato | Addgene | #130909 |
| **Experimental Models: Organisms/Strains** | | |
| *Pnoc*-IRES-Cre (PNOC-Cre) | The Jackson Laboratory | #034278 |
| *Slc32a1*-IRES-Cre (VGAT-Cre) | The Jackson Laboratory | #016962 |
| *Slc17a6*-IRES-Cre (VGLUT2-Cre) | The Jackson Laboratory | #016963 |
| *Slc17a6*-IRES2-FlpO x *Pnoc*-IRES-Cre (VGLUT2-Flp x PNOC-Cre) | Bred in house | N/A |
| **Reagents** | | |
| 2-Methylthiazoline (2MT) | Tokyo Chemical Industry Co | M0285 |
| Peppermint oil | Sigma Aldrich | 77411 |

### Table S2: Viruses, injection coordinates, and implant types

| Experiment | Mouse Line | Virus | Amount | Coordinates |
|---|---|---|---|---|
| pnVTA-PNOC stress response (Figures 1–3) | PNOC-Cre | AAV-DJ-Ef1a-DIO-GCaMP6s | 300nL | AP -3.25, ML 1.6, DV -4.65 (15° angle) |
| pnVTA retrograde tracing (Supplementary Figure 2) | VGAT-Cre, VGLUT2-Cre | AAV2-retro-Ef1a-DIO-eYFP | 300nL | AP -3.25, ML 0.4, DV -4.65 |
| LH-GABA/Glut to pnVTA-PNOC electrophysiology (Figure 4) | PNOC-Cre | AAV5-CaMKII-hChR2-eYFP | 300nL | AP -1.25, ML ±1.0, DV - 5.35 |
| | | AAV5-Ef1a-DIO-mCherry | 300nL | AP -3.25, ML ±0.4, DV -4.65 |
| LH-GABA odor response (Supplementary Figure 3) | VGAT-Cre | AAVDJ-Ef1a-DIO-GCaMP6s | 300nL | AP -1.25, ML ±1.0, DV - 5.35 |
| LH-Glut TST optogenetics (Supplementary Figure 3) | VGLUT2-Flp x PNOC-Cre | AAV-CAG-FLP-ChrimsonR-tdTom | 300nL | AP -1.25, ML ±1.0, DV - 5.35 |

**Supplementary Methods**

*Fiber photometry*

531-Hz sinusoidal LED light (Thorlabs, LED light: M470F3; LED driver: DC4104) was bandpass filtered (470 ± 20nm, Doric, FMC4) to excite GCaMP6s, a 211-Hz sinusoidal LED light (Thorlabs, M405FP1; LED driver: DC4104) was bandpass filtered (405 ± 10nm, Doric, FMC4) to evoke $Ca^{2+}$-independent isosbestic control emission. LED intensities were measured at the tip of the optic fiber and adjusted to 30µW before each recording. GCaMP6s emission traveled back through the same optic fiber then was bandpass filtered, (525 ± 25nm, Doric, FMC4), detected by a photoreceiver (Doric, DFD_FOA_FC), and recorded by a real-time processor (TDT, RZ5P). For the ChrimsonR stimulation experiments, a 635nm laser was passed through the filter cube at 1mW intensity to deliver red light at the tip of the optic fiber [32].

*Fiber photometry analysis*

Custom MATLAB scripts were used to detrend decay from bleaching by fitting a $4^{th}$ degree polynomial function to the raw signal (470nm) and isosbestic (405nm) traces, then dividing by the resulting curve. Then signal was normalized by dividing an LLS fit of the isosbestic trace scaled to the signal. The processed traces were then down-sampled by a factor of 100, smoothed across a rolling 1s window, extracted in windows surrounding the onset of relevant behavioral events (tail lift, odor, shock, air puff, looming, open arm entry, center entry), z-scored relative to the mean and standard deviation of each event window, and then averaged.

*Electrophysiology*

Coronal brain slices were prepared at 250 µM on a vibrating Leica VT1000S microtome using standard procedures. Mice were anesthetized with Isoflurane, and transcardially perfused with an ice-cold, oxygenated cutting solution consisting of (in mM): 93 N-Methyl-D-glucamine (NMDG), 2.5 KCL, 20 HEPES, 30 NaHCO3, NaH2PO4, 10 MgSO4·7H20, 0.5 CaCl2·2H20, 25 glucose, 3

$Na^+$-pyruvate, 5 $Na^+$-ascorbate, and 5 N-acetylcysteine. Following collection of coronal sections, the brain slices were transferred to a 34°C chamber containing oxygenated cutting solution for a 10-minute recovery period. Slices were then transferred to a holding chamber consisting of (in mM) 92 NaCl, 2.5 KCl, 20 HEPES, 2 MgSO4·7H20, 1.2 NaH2PO4, 30NaHCO3, 2 CaCl2·2H20, 25 glucose, 3 Na-pyruvate, 5 Na-ascorbate, 5 N-acetylcysteine and were allowed to recover for ≥ 30 min. For recording, slices were perfused with oxygenated artificial cerebrospinal fluid (ACSF; 31-33°C; 300-303 milliosmoles) consisting of (in mM): 113 NaCl, 2.5 KCl, 1.2 MgSO4·7H20, 2.5 CaCl2·6H20, 1 NaH2PO4, 26 NaHCO3, 20 glucose, 3 $Na^+$-pyruvate, 1 $Na^+$-ascorbate, at a flow rate of 2-3ml/min. For recordings of inhibitory currents, 10 μM CNQX was added to the external solution.
